# Single-cell transcriptome provides novel insights into antler stem cells, a cell type capable of mammalian organ regeneration

**DOI:** 10.1101/263798

**Authors:** Hengxing Ba, Datao Wang, Weiyao Wu, Hongmei Sun, Chunyi Li

**Affiliations:** State Key Laboratory for Molecular Biology of Special Wild Economic Animals, Institute of Special Wild Economic Animals and Plants, Chinese Academy of Agricultural Sciences, Changchun 130112, China; BGI Genomics, BGI-Shenzhen, Shenzhen 518083, China.; Changchun Sci-Tech University

**Keywords:** antler, stem cell, antlerogenic periosteum, single cell, transcriptome, scRNA-seq

## Abstract

Antler regeneration, a stem cell-based epimorphic process, has potential as a valuable model for regenerative medicine. A pool of antler stem cells (ASCs) for antler development is located in the antlerogenic periosteum (AP). However, whether this ASC pool is homogenous or heterogeneous has not been fully evaluated. In this study, we produced a comprehensive transcriptome dataset at the single-cell level for the ASCs based on the 10x Genomics platform (scRNA-seq). A total of 4,565 ASCs were sequenced and classified into a large cell cluster, indicating that the ASCs resident in the AP are likely to be a homogeneous population. The scRNA-seq data revealed that tumor-related genes were highly expressed in these homogeneous ASCs: i.e. TIMP1, TMSB10, LGALS1, FTH1, VIM, LOC110126017 and S100A4. Results of screening for stem cell markers suggest that the ASCs may be considered as a special type of stem cell between embryonic (CD9) and adult (CD29, CD90, NPM1 and VIM) stem cells. Our results provide the first comprehensive transcriptome analysis at the single-cell level for the ASCs, and identified only one major cell type resident in the AP and some key stem cell genes, which may hold the key to why antlers, the unique mammalian organ, can fully regenerate once lost.

## Introduction

The ‘Holy Grail’ of modern regenerative medicine is to grow back lost organs or appendages, which is known as epimorphic regeneration (RJ 1983; Stocum 2006). Our current knowledge of epimorphic regeneration is largely gained from the studies on lower vertebrates (Gardiner et al. 2002). Notably, these animals have the ability to reprogram phenotypically-committed cells at the amputation plane toward an embryonic-like cell phenotype (de-differentiation) and to form a cone-shaped tissue mass, known as a blastema (Mescher 1996). Deer antlers are the only mammalian appendages capable of full renewal and therefore offer a unique opportunity to explore how nature has solved the problem of epimorphic regeneration in mammals (Goss 1995; Kierdorf et al. 2009; Li et al. 2009; Li 2012). Recent studies concluded that antler regeneration is a stem cell-based epimorphic process (Kierdorf et al. 2007; Li et al. 2005; Li et al. 2007a; Rolf et al. 2008), and has the potential for development as a valuable model for biomedical research and regenerative medicine. Revealing the mechanism underlying this stem cell-based epimorphic regeneration in mammals would undoubtedly place us in a better position to promote tissue/organ regeneration in humans.

Antlers regenerate from the permanent cranial bony protuberances, known as pedicles. Growth of a pedicle itself is initiated when a male deer approaches puberty. The origin is a piece of periosteum, known as antlerogenic periosteum (AP), which covers the frontal crest on the skull (Li and Suttie 1994). Removal of the AP prior to pedicle initiation stops pedicle and antler growth, and transplantation of the AP autologously induces ectopic pedicle and antler formation (Goss and Powel 1985; Li et al. 2002; Li et al. 2007b). The initial discovery of the AP (H and J 1974) has been hailed as “hallmark” event in antler research history (RJ 1983). The AP tissue, ~2.5 cm in diameter and 2.5-3 mm in thickness, contains around five million cells, which sustain the seasonal renewal of the entire antlers throughout the deer’s life (Li et al. 2009). Potency of the AP cells has been investigated by several laboratories (Berg et al. 2007; Li and Suttie 2001; Price et al. 2005a; Rolf et al. 2008). The AP cells can be induced *in vitro* to differentiate into chondrocytes, osteoblasts, adipocytes, myoblasts and neural-like cells. Therefore, AP cells have been termed antler stem cells (ASCs) and are essential for full regeneration of this unique mammalian organ (Li et al. 2009).

Differences in cell type within any tissue are essential for their biological states and function. Numerous studies in cell biology have utilized single-cell sequencing by employing new protocols of single cell isolation to characterize functionally heterogeneous cells (Yu and Lin 2016). This study is the first to apply single-cell sequencing technology to investigate the ASCs through transcriptome (scRNA-seq) using the 10x genomics platform, a droplet-based system that enables 3’ messenger RNA (mRNA) digital counting for thousands of single cells (Zheng et al. 2017).

## Materials and Methods

### AP tissue sampling

The AP tissues were obtained from a 6-month-old male sika deer immediately after slaughtering, according to the previous protocol (Li and Suttie 2003). Briefly, to collect the AP tissue, a crescent-shaped incision was made on the scalp skin 2 cm medial to the frontal crest; skin was separated from the frontal bone to expose the AP. The AP was then peeled from the underlying bone following the delineating incisions cut on the periosteum and then placed into 50 ml centrifuge tube containing 20 ml cold DMEM (Gibco; Grand Island, USA) plus 500 U/ml penicillin and 500 g/ml streptomycin (Invitrogen, USA). All experimental processes were approved by Animal Ethics Committee of Institute of Special Wild Economic Animals and Plants, Chinese Academy of Agricultural Sciences (CAAS2017015).

### Isolation of the AP cells

Isolation of the AP cells was carried out according to our previous methodology (Li et al. 2012; Li et al. 1999). Briefly, after sampling, AP tissue was immediately cut into thin slices (around 0.7 mm in thickness) using a custom-built tissue cutter (Chu et al. 2017). These slices were digested in the DMEM culture medium containing collagenase (150 units/ml) at 37°C for 1-1.5 hour to release cells, and the released cells were cultured in a medium (DMEM +10% FBS +100 U/ml penicillin +100 mg/ml streptomycin). In order to increase cell numbers to meet the requirement for subsequent construction of the scRNA-seq library, the cells were trypsinized when reaching confluence and then reseeded in T75 culture flasks at a density of 5×10^5^ cells/ml for one more round of expansion.

### Single cell sequencing using ChromiumTM Platform

The scRNA-seq library was constructed using the Chromium™ Controller and Chromium™ Single Cell 3’ Reagent Version 1 Kit (10x Genomics, Pleasanton, CA) to generate single cell GEMs (Gel Bead-in-Emulsions) as previously described (Zheng et al. 2017). Briefly, about 5×10^5^/ml (500/μl) suspended cells was obtained and placed on the ice. In total, a 15 μl cellular suspension that contained ~7500 cells was added to the Master Mix in the tube strip well. The 100 μl Master Mix containing cells, 40 μl Single Cell 3’ Gel Beads and 135 μl oil-surfactant solution were transferred to each well in the ChromiumTM Single Cell 3’ chip row. Subsequently, GEM-RT was performed using a Thermocycler (BioRad; 55 °C for 2 hours, 85 °C for 5 mins, hold at 4 °C). Post GEM-RT, cleanup and cDNA amplification was performed to isolate and amplify cDNA for library construction. The samples were sequenced in two lanes on the HiSeq 2500 in rapid run mode using a paired end flow cell. Read1: 98 cycles, Index1: 14 cycles, Index2: 8 cycles, and Read2: 10 cycles.

### scRNA-seq data analysis

Cell Ranger Software Suite version 1.3.1 (http://support.10xgenomics.com/) was used to perform sample de-multiplexing, barcode processing and single-cell 3′ gene counting, as performed previously (Zheng et al. 2017). Ten bp transcripts/Unique Molecular Identifier (UMI) tags were extracted from Read2. Cellranger mkfastq used bcl2fastq v2.19 (https://support.illumina.com/) to demultiplex raw base call files from Hiseq2500 sequencer into sample-specific FASTQ files. Cellranger mkref was run to produce a cellranger-compatible reference based on both the Ovir.te1.0 genome sequences and the transcriptome GTF file. These FASTQ files were aligned to the reference with cellranger count that used an aligner called STAR (Dobin et al. 2013).

The cells were selected based on the following criteria (Yan et al. 2017): 1) number of expressed genes (300-5000); 2) number of UMI counts (<25,000); 3) percentage of mitochondrial genes (<5%); and 4) number of cells expressed per gene (≥5). After normalizing expressed data by NormalizeData function, dispersion of each gene against the mean expression level was plotted using FindVariableGenes function (x.low.cutoff = 0.01, x.high.cutoff = 4, y.cutoff = 0.3), which was in Seurat R package version 2.2.1 (Butler et al. 2018). A total of 2,943 variable genes were selected based on their plotting results (Fig. S1) and the amount of variability was found to be explained by cell cycle genes. The cell cycle scores were further generated by CellCycleScoring function based on G2/M and S phase markers, and then, these cores were used to scale cell-gene expression data using ScaleData function. The standard deviations of the principle components were plotted by PCElbowPlot function to identify the true dimensionality of a dataset (Fig. S2). Based on unsupervised graph-based nearest neighbor clustering algorithm with different resolution degrees, the scaled expression datasets were clustered using FindClusters function with the first 30 principal components according to the principal components analysis elbow plot. The cluster results were presented using t-distributed stochastic neighbor embedding (t-SNE)-based plots (Blondel et al. 2008). The FindAllMarkers function was used to find differentially expressed genes with absolute log2 foldchange >1 and adjust P-value < 0.001.

### Immunofluorescent staining

The primary cultured ASCs were fixed with 4% paraformaldehyde for 30 minutes and blocked by incubation in 3% BSA (0.1% Triton X-100 for NPM1) in PBS for 1 hour at RT, followed by incubation with the primary antibodies overnight at 4 °C. The cells were incubated with secondary antibody for 30 min followed by DAPI (blue) staining for visualization of nuclei. The primary and secondary antibodies used in the study are listed in Table S1. The primary antibodies were replaced by rabbit or mouse IgG for the isotype-matched controls. All images were captured under a fluorescent microscope (EVOS, ThermoFisher, USA).

### Flow cytometry analysis

The ASCs were incubated with the primary antibodies overnight at 4 °C, and then the cells were stained with FITC-conjugated secondary antibodies for another 1 hour at RT. Isotype-matched rabbit or mouse IgG was used as a negative control. After 3 times of washes with cold PBS, the cells were resuspended in 500 μl PBS. Flow cytometry analysis was performed using FACSCalibur and the results were analyzed using Cellquest software (BD Biosciences; USA).

## Results

### High-quality scRNA-seq data

A total of 252,818,309 read pairs were mapped to 14,993 genes, which were found to be expressed in the 4,731 sorted individual ASCs. This was equivalent to an average of 53,438 mapped read pairs per cell, which is reportedly sufficient for an accurate analysis by Single Cell 3′ Solution (Yan et al. 2017). The median gene number and UMI counts were 2,568 and 10,309 respectively. The results of detailed statistical analysis of the scRNA-seq data and the sequenced cells are summarized in Table S2 and Table S3 respectively.

A steep drop-off of barcode UMI counts was indicative of good separation of the cell-associated barcodes from the empty droplets (Fig. 1A). The number of genes (Fig. 1B) and percentage of mitochondrial genes (Fig. 1C) against the corresponding UMI counts per cell were plotted respectively to exclude outlier cells as potential multi-cell-droplets. Based on the threshold criteria (see Methods), we filtered out 116 cells and 1,820 genes. Altogether, 4,615 singular cells and 13,173 genes were retained for further analysis.

**Figure 1.**
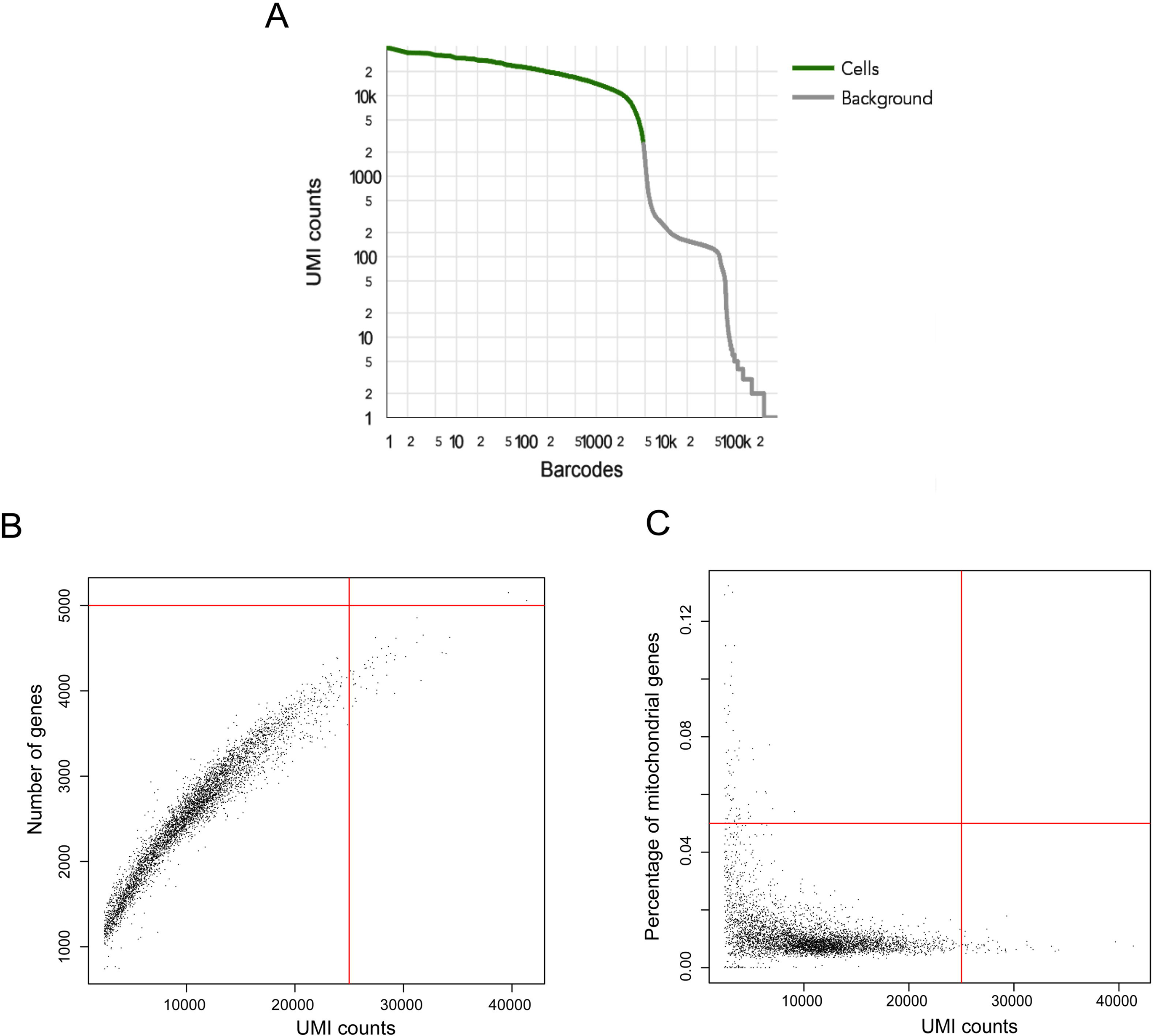
Quality metrics of the ASC single-cell transcriptomes using 10x Genomics. **A)** Barcode rank plot. In the plot, a steep drop-off is indicative of good separation between the cell-associated barcodes and the barcodes associated with empty partitions. B) Plot between number of genes and UMIs counts per cell. C) Plot of mitochondria UMIs and UMIs counts per cell. Cells (4,615 in total) and UMIs (25,000) were selected for downstream analysis (red dashed lines). Cells with number of genes <5,000 and >300, UMI counts <25,000, percentage of mitochondrial genes <5% were selected for downstream analysis (red dashed lines).

### A large cell cluster across the ASCs

The graph-based nearest-neighbor clustering algorithm that did not rely on known markers uncovered two cell clusters from 4,615 high-quality cells with a resolution degree of 0.1 (Fig. 2A). We further defined groups of genes (absolute log2 foldchange >1 and adjust P-value < 0.001), which allowed classification of these cells into two distinct cell clusters (Fig. 2B). We found no up-regulated genes in the small cluster (50 cells). However, a high proportion of down-regulated mitochondrial genes (i.e., MT-COX1, MT-COX2, MT-COX3, MT-ND2, MT-ND1 and MT-CYTB) were represented, strongly suggesting that the small cluster could be contaminated by debris of the ASCs. We also found the ASCs in the large cell cluster (4,565 cells) expressed high levels of extracellular matrix proteins, such as collagen family. In particular, these genes (i.e. COL1A1, COL1A2, COL5A1, COL5A2, SPARC and FN1) were typically applied to measure differentiated states from mesenchymal stem cell to osteoblast, osteoprogenitor and chondrogenesis. Based on the resolution degree of 0.2, the large cell cluster was again separated into two clusters with 2,404 and 2,161 cells respectively (Fig. 2C), but only four genes had values of absolute log2 foldchange >1 and <2 (ACTA2, THBS1, TNC and MYL9) (Fig. 2D), indicating that the ASCs resident in the AP are likely to represent a homogeneous population.

**Figure 2.**
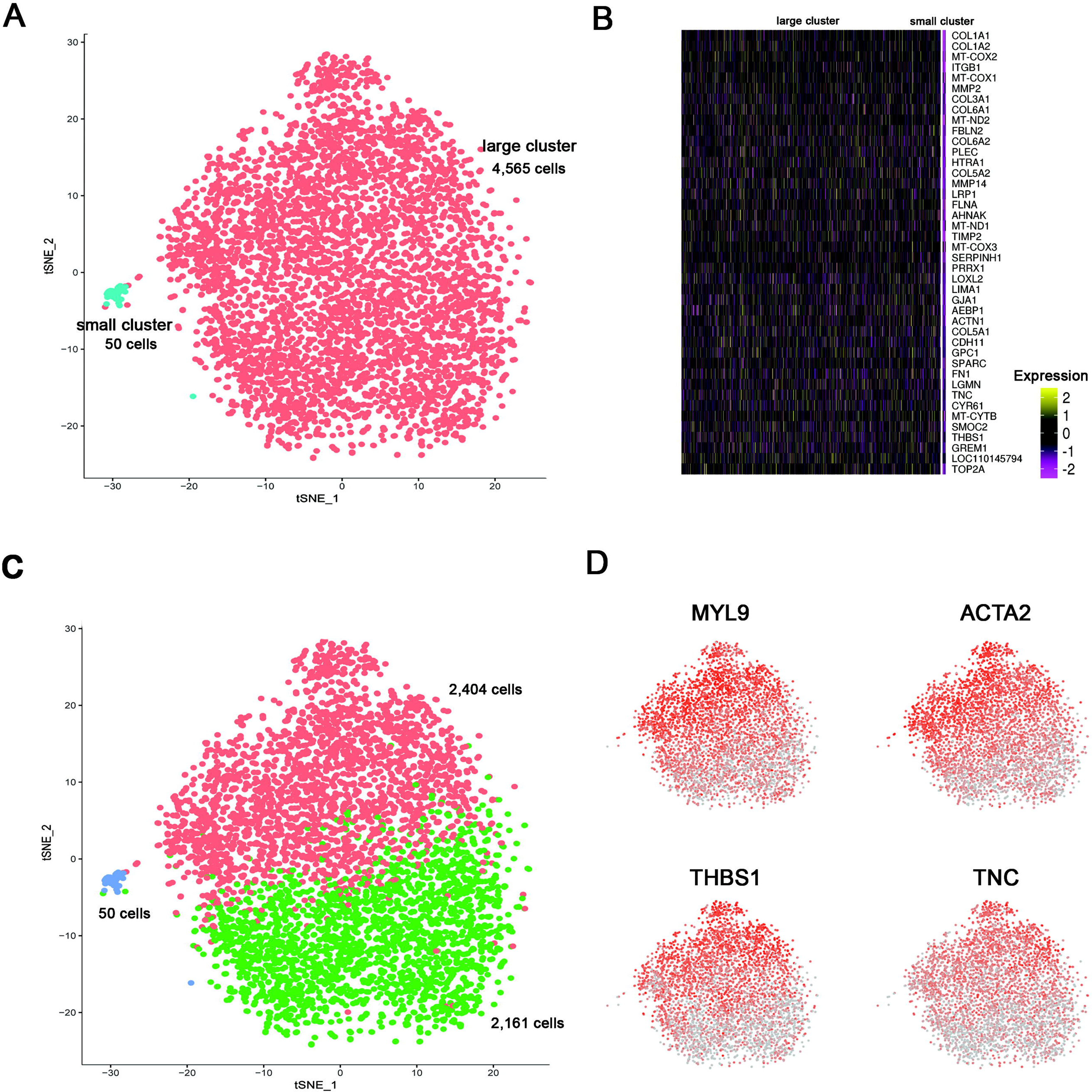
Unsupervised graph based nearest neighbor clustering analysis of the ASCs. A) t-SNE projection of single cells, colored by two inferred ASC clusters based on the resolution degree of 0.1. B) Normalized expression (centred) of differentially expressed genes (rows) from each of two clusters (columns) is shown in a heatmap. Gene symbols are represented at the right. C) t-SNE projection of single cells, colored by three inferred ASC clusters based on the resolution degree of 0.2. D) Four genes that have values of absolute log2 foldchange >1 and <2 are labeled on t-SNE plots.

### Highly expressed genes across the ASCs

A total of 35 genes were found to be highly and commonly expressed in cells of the large cell cluster (at least 20 UMIs); these were further sorted based on their expression levels (Fig. 3A and Table S4). Of those selected 35 genes, the first seven, i.e., Metalloproteinase inhibitor 1 (TIMP1), Thymosin beta 10 (TMSB10), Galectin-1 (LGALS1), Ferritin heavy chain (FTH1), Vimentin (VIM), ferritin light chain-like (LOC110126017) and (S100A4, were found to be expressed over 50 UMIs. Almost all of the ASCs expressed these genes above the level of 10 UMIs. Notably, 98% and 90% of the ASCs expressed TMSB10 gene above the level of 30 and 50 UMIs respectively. The expression levels of the TMSB10, LGALS1 (Fig. 3B) and VIM genes (refer to Fig. 4A) were further confirmed using immunofluorescent staining. Of the selected 35 genes, 25 (71%) were found to be involved in the protein-protein interaction network in the STRING v10.5 (Fig. 3C), suggesting that they may come together to accomplish a particular biological function within the ASCs.

**Figure 3.**
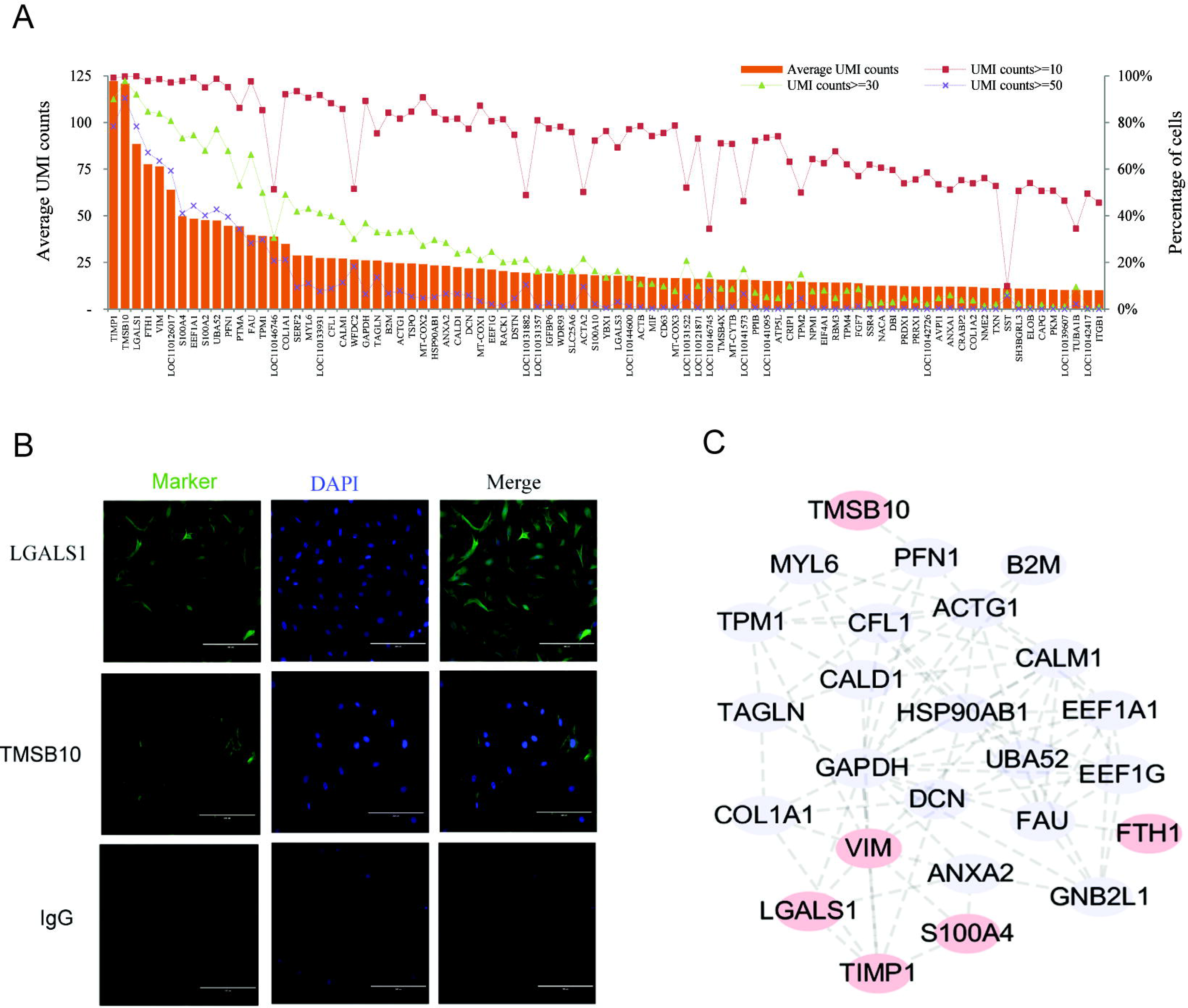
Highly expressed genes in the scRNA-seq data. A) Top 35 genes that were expressed ≥ 20 UMIs averaged across all the ASCs (bar plot). Percentage of cells based on ≥ 10, 30 and 50 UMIs per cell respectively (line plot). B) Immunostaining of the ASCs using anti-TMSB10 and anti-LGALS1. DAPI staining was used for nuclei detection. IgG staining was used as negative control. Scale bar: 200 m. C) Protein-protein interaction network between 25 of the top 35 genes based on STRING v10.5 (string-db.org) searching with medium confidence (0.4).

**Figure 4.**
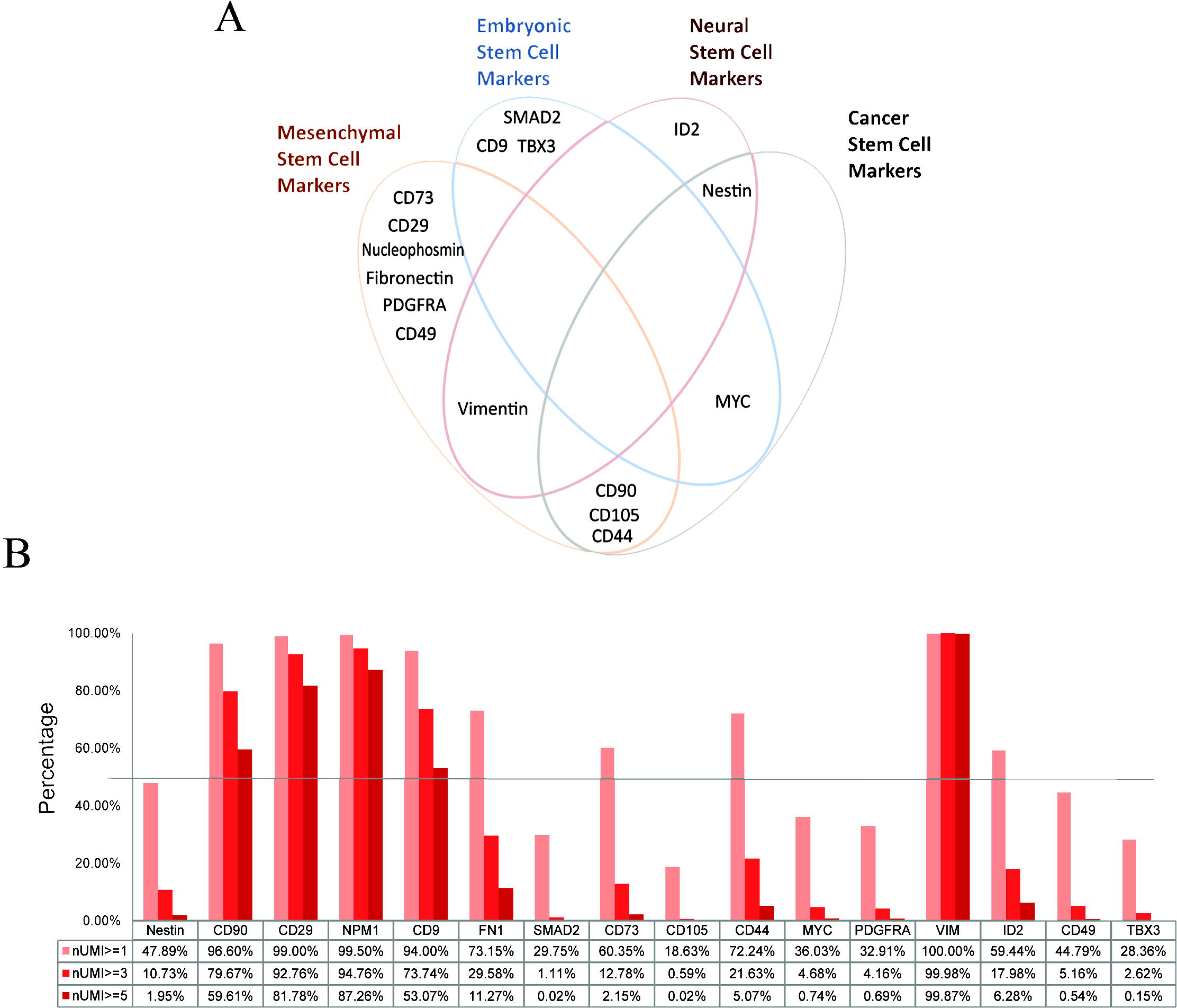
ASC screening results using the currently available stem cell markers. These cells were labeled by 16 stem cell markers respectively, and the label threshold was set to meet a criterion that a marker must be expressed by more than 3% ASCs and with at least one UMI count. A) Venn diagram of the 16 individual stem cell markers across the four types of stem cell marker, including 10 mesenchymal stem cell markers, four embryonic stem cell markers, three neural stem cell markers and five cancer stem cell markers. B) Expression abundance of the 16 individual stem cell markers based on UMI counts ≥ 1, 3 and 5.

### Expression of stem cell markers

In order to investigate expression status of stem cell markers for the ASCs in the large cell cluster, we selected 16 marker genes from the scRNA-seq data based on two criteria: 1) expressed in at least one UMI count; 2) expressed in over 3% of the ASCs. These 16 marker genes were then classified into four types of stem cell markers based on the definition for each type from the literature: 1) mesenchymal stem cell markers (10; CD29, CD73, CD105, CD90, Fibronectin (FN1), VIM, Nucleophosmin (NPM1), PDGFRA, CD44 and CD49); 2) embryonic stem cell markers (4; CD9, SMAD2, MYC and TBX3); 3) neural stem cell markers (3; ID2, Nestin (NES) and VIM); and 4) cancer stem cell markers (4; CD44. MYC, CD90 and CD105) (Fig. 4A). When the expression level was set at nUMI > 5, four mesenchymal (CD90, CD29, VIM and NPM1) and one embryonic stem cell marker genes (CD9) were found to be expressed in over 50% of the ASCs (Fig. 4B). More significantly, VIM, known as a neural stem cell marker (Bramanti et al. 2010), was highly expressed in almost all of the ASCs.

High expression levels for these five stem cell marker genes (CD9, CD90, CD29, VIM and NPM1) were further confirmed using immunofluorescent staining (Fig. 5A). In addition, the immunofluorescent staining results provided extra information over the expression level study. For example, one of the NPM1 functions is to act on genomic stability and DNA repair (Lindstrom 2011), and in our results distinct dots were clearly evident within and around the nucleus of the ASCs, suggesting that it is a nucleus protein. Also as expected, VIM, a major cytoskeletal protein of mesenchymal cells (Colucci-Guyon et al. 1994; Goldman et al. 1996), was observed to be in a distinct form outside of nuclei. To further investigate the proportion of positive cells for each of these five marker genes, FACS analysis was performed and the results showed that CD9^+^, CD90^+^, CD29^+^, VIM^+^ and NPM1^+^ were positive in 91.1%, 93.2%, 92.8%, 98.4% and 94.5% of ASCs respectively (Fig. 5B), which were concordant with the results of scRNA-seq data analysis.

**Figure 5.**
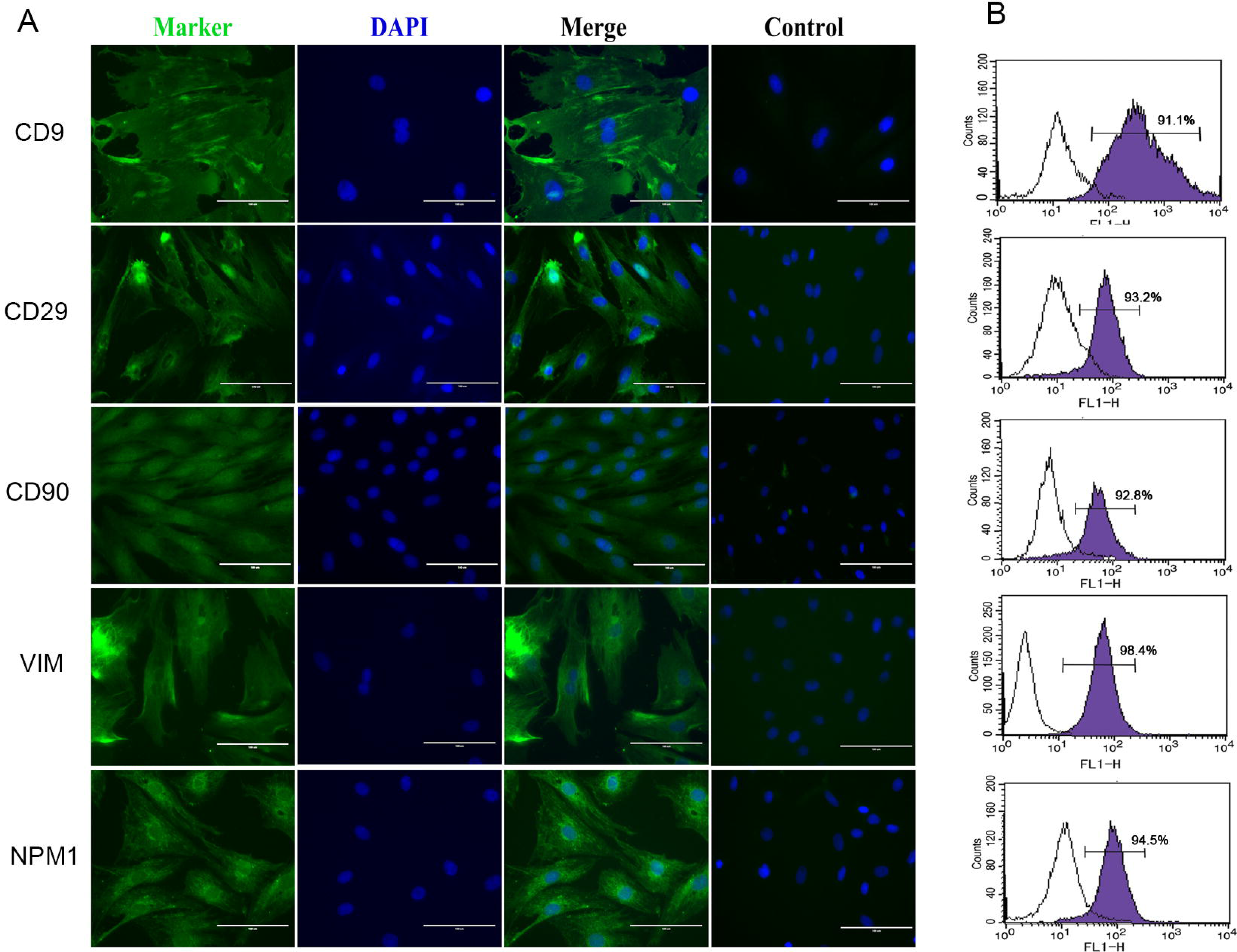
Immunostaining and FACS analysis of the ASCs. A) Immunostaining of ASCs using anti-CD9, anti-CD29, anti-CD90, anti-VIM and anti-NPM1 antibodies. DAPI staining was used for detection of nuclei. Scale bar: 200μm. B) FACS analysis was performed using each of these five antibodies. Values show the intensity of the indicated antigen.

## Discussion

Single-cell RNA sequencing (scRNA-seq) approaches have become increasingly popular providing insights into various aspects of developmental and stem cell biology (Kumar et al. 2017). In the present study, using the scRNA-seq technology based on the 10x genomics platform, we provided an initial high-resolution picture of molecular characterization for the ASCs (4,615 cells including 13,173 genes).

Several highly expressed genes, such as S100A4, LGALS1 and TMSB10, have been reported previously as being related to the development of the AP tissues/cells (Li et al. 2012; Park et al. 2004; Wang et al. 2017), and TMSB10 is also highly expressed in growing antlers (Lord et al. 2004; Zhang et al. 2018). These highly-expressed genes are reportedly tumor-related factors. S100A4 promotes cell proliferation and tumor growth (Sherbet 2009). The down-regulation of S100A4 expression suppresses cell proliferation in many cancer cells (Huang et al. 2012; Ma et al. 2010). LGALS1 modulates the immune response (Liu 2005; Rabinovich et al. 2007) and may contribute to immune privilege in tumors. TMSB10 plays important roles in the progression and metastasis of various tumors (Santelli et al. 1999; Sribenja et al. 2009; Zhang et al. 2017). The dysregulated activity of TIMP1 has been implicated in tumors (Kim et al. 2012). Methylation of VIM has been established as a biomarker of tumors (Jung et al. 2011). Both FTH1 and LOC110126017 (ferritin light chain-like) encode proteins that play important roles in iron storage and homeostasis. It is known that iron has a role in the tumour microenvironment and in metastasis and can contribute to both tumour initiation and tumour growth (Torti and Torti 2013). Although these tumor-related genes are highly expressed in the ASCs, these cells do not become cancerous during antler formation despite an astonishing rate of proliferation and differentiation, which is genuinely impressive and worth further exploration.

Recently, it has been reported that the ASCs express both mesenchymal stem cell markers, such as STRO-1 and CD105, and embryonic stem cell markers such as CD9 and MYC (Li et al. 2009; Rolf et al. 2008; Seo et al. 2014). Surprisingly, the ASCs were also reported to express some key embryonic stem cell marker genes, such as Oct4, SOX2 and NANOG (Li et al. 2009; Seo et al. 2014). However, we failed to detect the expression of these key embryonic marker genes through the scRNA-seq in this study. Interestingly, expressed NANOG in the ASCs was found as a pseudogene in one of our previous studies (Wang et al. 2016). In addition, the ASCs have be considered to be of neural-crest origin (Kierdorf et al. 2007; Li and Suttie 2001; Price et al. 2005b), with direct evidence being the detection of the mRNAs for several neural crest-cell markers in the ASCs using RT-PCR (Mount et al. 2006). The neural crest cell population is an embryonic cell population and these cells might represent some kind of “embryonic remnant” comprising pluripotent cells left over from the early embryo (Li et al. 2009). Our results also showed that almost all of the ASCs highly expressed the VIM gene, a marker that appears at the earliest stage of neural tube development (Houle and Fedoroff 1983), which further supports the neural crest origin of the AP cells. Overall, despite the lack of expression of key embryonic marker genes in this study, the ASC may still be considered as a unique type of stem cell that has biological attributes derived from embryonic (CD9), mesenchymal (CD29, CD90, NPM1 and VIM) and neural stem cells (VIM). Our findings in this study strongly support the view that the annual antler regeneration represents a stem cell-based process.

The scRNA-seq studies have thus far led to the discovery of novel cell types and provided insights into regulatory networks during development (Liu and Trapnell 2016). Using this powerful approach, we have successfully identified only one major type of ASC resident in the AP. It is understandable that, for such a small piece of tissue (around 2.5 cm in diameter and 2 mm in thickness) to initiate a large mammalian appendage (up to 15 kg) within two to three months of time, this limited number of cells (around 5 million) must uniformly possess stem cell attributes, such as almost unlimited proliferation potential. Whatever it is, the results from the present study provide a useful source for further investigation at molecular level of deer antler renewal, the only stem cell-based mammalian organ regeneration.

## Supporting information

Detection of variable genes across the ASCs

Standard deviation of principal components

List of Antibodies

Summary of scRNA-seq data quality

Summary of single cell sequencing data

Summary of top 35 highly expressed genes in the scRNA-seq data

## Availability of supporting data

The raw single-cell RNA-seq data in fastq format can be found at SRA under BioProject PRJNA416396.

## Acknowledgements

We wish to thank Drs. Peter Fennessy and Eric Lord for reading through the paper and giving valuable comments.

## Author contributions

H.B., D.W., and C.L. conceived the experiment. D.W. collected the samples and performed molecular- and cell-related experiments. H.S. cultured the cell lines. W.W. extracted the RNA samples and prepared them for single-cell 3′ library construction and sequencing. H.B. performed QC and data analysis. H.B., W.W., and C.L. wrote the manuscript. All authors read and approved the final manuscript.

---

**Figure S1. Detection of variable genes across the ASCs.** A total of 2,415 variable genes were selected by using FindVariableGenes function (x.low.cutoff = 0.01, x.high.cutoff = 4, y.cutoff = 0.3) in Seurat R package. The parameters identify ~2,943 variable genes.

**Figure S2**. **Standard deviation of principal components.** To assess the true dimensionality of our dataset, the first 30 principal components were deemed as a cutoff, as there is a clear elbow in the graph.

Table S1. List of Antibodies

Table S2. Summary of scRNA-seq data quality Table S3. Summary of single cell sequencing data

Table S4. Summary of top 35 highly expressed genes in the scRNA-seq data.

