## Supplementary figures and images for "Single-cell transcriptome provides novel insights into antler stem cells, a cell type capable of mammalian organ regeneration"

### Detection of variable genes across the ASCs

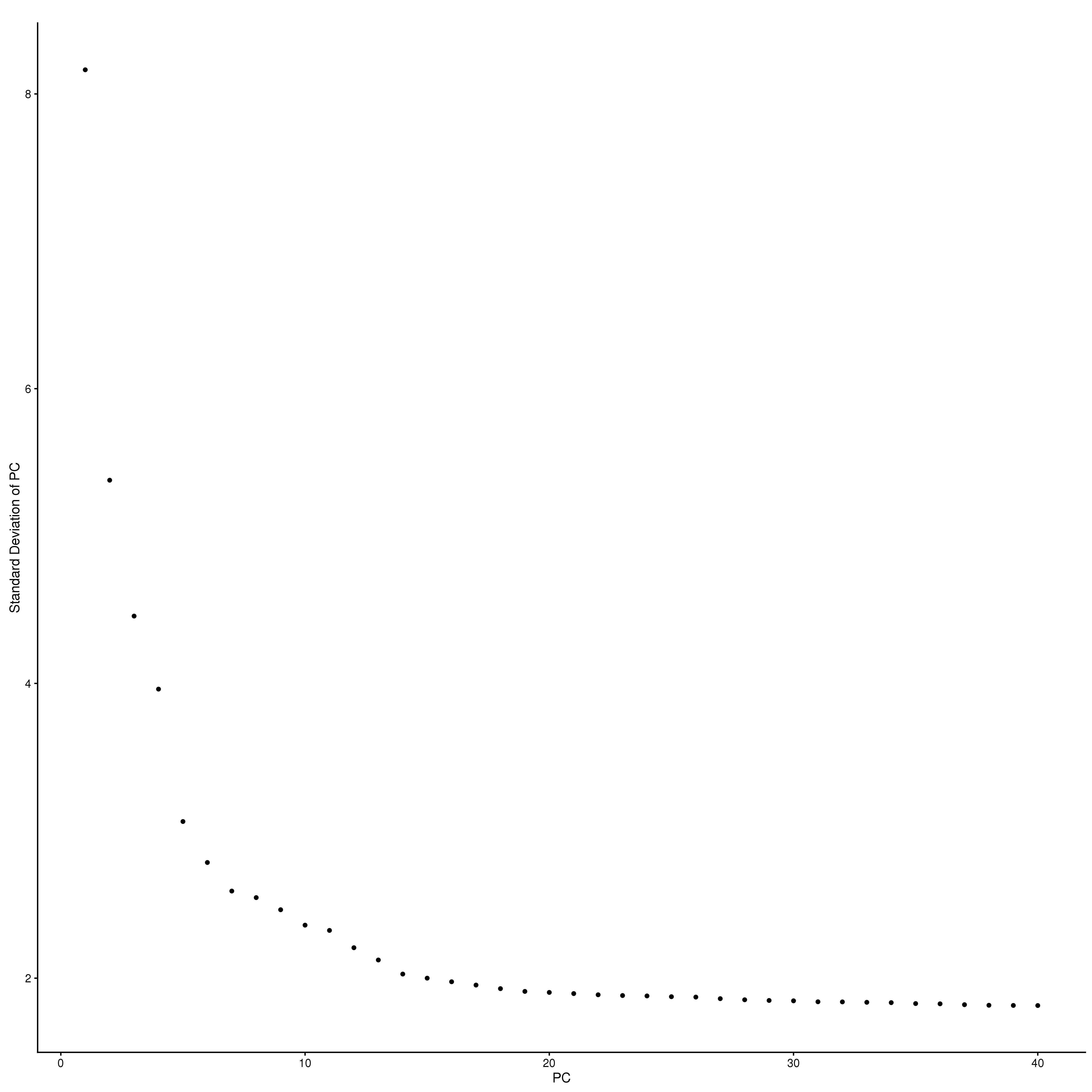

### Standard deviation of principal components

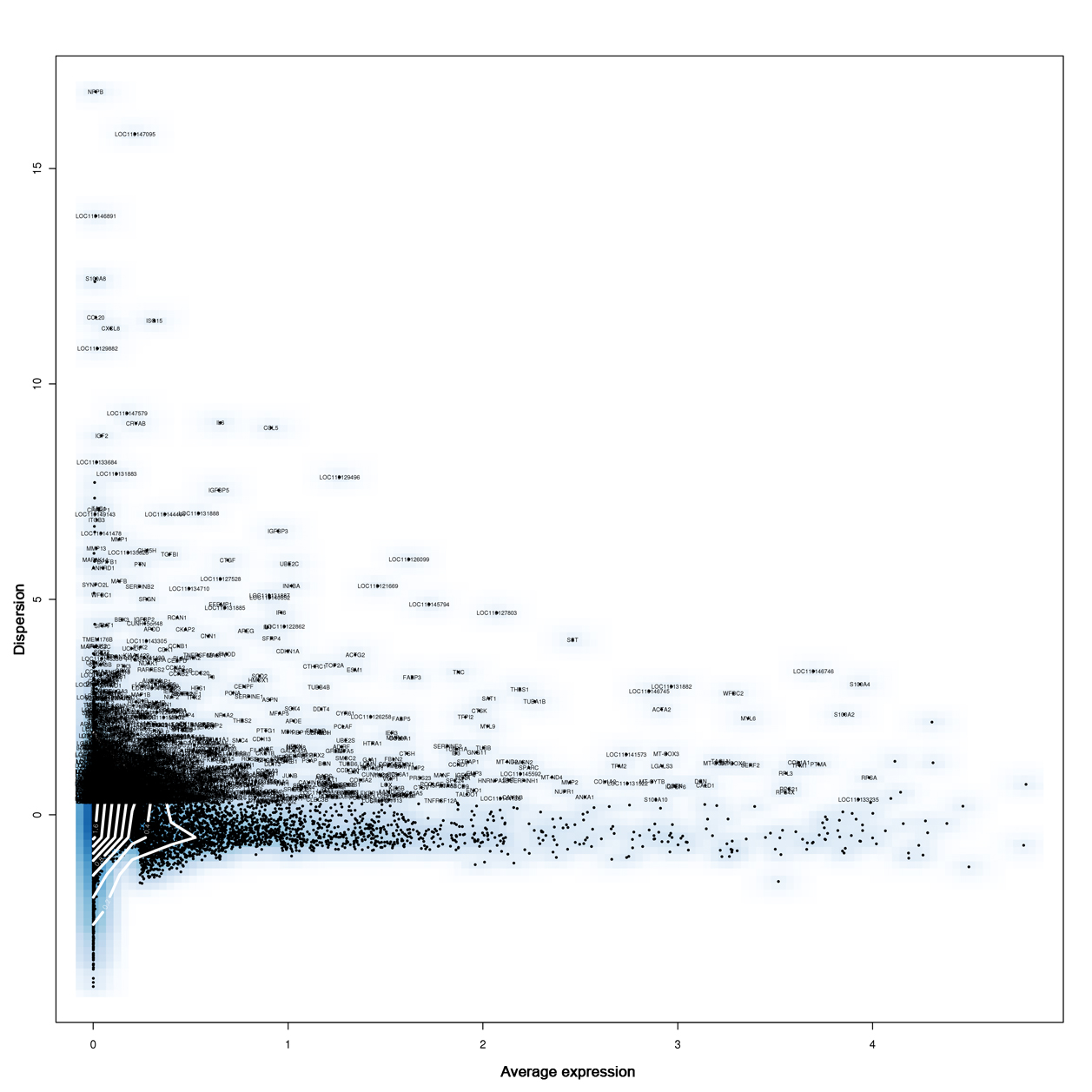
