## Supplementary material for "Single-cell transcriptome provides novel insights into antler stem cells, a cell type capable of mammalian organ regeneration": List of Antibodies

Table S1. List of Antibodies

| **Terms** | **Source** | **Type** | **Company** | **Catalog number** | **Dilution** |
| --- | --- | --- | --- | --- | --- |
| Thymosin beta 10 (TMSB10) | Rabbit | Polyclonal | Abcam, USA | Ab14338 | IF:1:100, FLC:1:200 |
| Galectin 1 (LGALS1) | Rabbit | Polyclonal | Bioss, China | Bs-6594R | IF:1:100, FLC:1:200 |
| CD9 | Mouse | Monoclonal | ThermoFisher, USA | MA1-19301 | IF:1:100, FLC:1:200 |
| CD29 | Mouse | Monoclonal | R&D Systems, USA | MAB17783 | IF:1: 100, FLC:1:200 |
| CD90 | Mouse | Monoclonal | R&D Systems, USA | MAB2067 | IF:1:100, FLC:1: 200 |
| Vimentin (VIM) | Mouse | Monoclonal | Abcam, USA | Ab8978 | IF:1:100, FLC:1:200 |
| Nucleophosmin 1 (NPM1) | Rabbit | Polyclonal | Bioss, China | Bs-4757R | IF:1:100, FLC:1:200 |
| IgG-Isotype control | Rabbit |  | Abcam, USA | Ab172730 | IF:1:100, FLC:1:200 |

FLC: flow cytometry; IF: immunofluorescence
