## Supplementary material for "Single-cell transcriptome provides novel insights into antler stem cells, a cell type capable of mammalian organ regeneration": Summary of scRNA-seq data quality

Table S2. Summary of single cell sequencing quality

| **Sequencing quality metrics** | **Value** |
| --- | --- |
| Number of Reads | 252,818,309 |
| Valid Barcodes | 94.7% |
| Reads Mapped Confidently to Transcriptome | 61.3% |
| Reads Mapped Confidently to Exonic Regions | 63.5% |
| Reads Mapped Confidently to Intronic Regions | 5.4% |
| Reads Mapped Confidently to Intergenic Regions | 8.8% |
| Sequencing Saturation | 58.5% |
| Q30 Bases in Barcode | 72.9% |
| Q30 Bases in RNA Read | 93.6% |
| Q30 Bases in Sample Index | 96.8% |
| Q30 Bases in UMI | 97.0% |
