## Supplementary material for "Single-cell transcriptome provides novel insights into antler stem cells, a cell type capable of mammalian organ regeneration": Summary of single cell sequencing data

Table S3. Summary of single cell sequencing cells

| **Sequencing cells metrics** | **Value** |
| --- | --- |
| Estimated Number of Cells | 4,731 |
| Fraction Reads in Cells | 82.9% |
| Mean Reads per Cell | 53,438 |
| Median Genes per Cell | 2,568 |
| Total Genes Detected | 14,993 |
| Median UMI Counts per Cell | 10,309 |
